# Transsaccadic integration operates independently in different feature dimensions

**DOI:** 10.1101/2021.06.03.446892

**Authors:** Garry Kong, David Aagten-Murphy, Jessica M. V. McMaster, Paul M. Bays

**Affiliations:** Department of Psychology, University of Cambridge, United Kingdom

## Abstract

Our knowledge about objects in our environment reflects an integration of current visual input with information from preceding gaze fixations. Such a mechanism may reduce uncertainty, but requires the visual system to determine which information obtained in different fixations should be combined or kept separate. To investigate the basis of this decision, we conducted three experiments. Participants viewed a stimulus in their peripheral vision, then made a saccade that shifted the object into the opposite hemifield. During the saccade, the object underwent changes of varying magnitude in two feature dimensions (Experiment 1: color and location, Experiments 2 and 3: color and orientation). Participants reported whether they detected any change and estimated one of the post-saccadic features. Integration of pre-saccadic with post-saccadic input was observed as a bias in estimates towards the pre-saccadic feature value. In all experiments, pre-saccadic bias weakened as the magnitude of the transsaccadic change in the estimated feature increased. Changes in the other feature, despite having a similar probability of detection, had no effect on integration. Results were quantitatively captured by an observer model where the decision whether to integrate information from sequential fixations is made independently for each feature and coupled to awareness of a feature change.

## Introduction

With each saccadic eye movement, incoming visual information undergoes an abrupt transformation. Not only does the shift in gaze direction create a substantially different image on the retina, but the transition between these snapshots is obscured by blur induced by the movement itself. Our smooth and continuous perceptual experience during this process makes it easy to overlook the complex computations required by the visual system to coordinate it. According to contemporary theories, a limited amount of visual information from before a saccade, primarily from the region of the intended target, is maintained and used to resolve perception after the saccade (Burr & Morrone, 2011; Cavanagh, Hunt, Afraz, & Rolfs, 2010; Melcher & Morrone, 2015), while changes outside this focus go unnoticed due to limitations of attention and working memory (Aagten-Murphy & Bays, 2018; Rensink, O’Regan, & Clark, 1997).

One line of evidence for these theories has come from studies investigating the phenomenon of Suppression of Saccadic Displacement (Bridgeman, Hendry, & Stark, 1975), whereby observers may fail to notice large shifts of a saccade target occurring during the eye movement (as much as half the saccade amplitude). This has been explained on the basis that the eye movement itself creates a large and uncertain shift of object location on the retina, making it difficult to determine whether to attribute retinal displacements to the saccade or to a change in object location (Niemeier, Crawford, & Tweed, 2003). Supporting this account, factors that would favor the hypothesis that the object has changed aid in detection of a displacement, e.g., an intrasaccadic change to a surface feature of the object, or a brief blank interval before reappearance of the object after the saccade (Demeyer, De Graef, Wagemans, & Verfaillie, 2010; Poth, Herwig, & Schneider, 2015; Tas, Moore, & Hollingworth, 2012; Wexler & Collins, 2014).

There is, however, even more direct evidence that pre-saccadic input is available and used by the visual system after the saccade. When the surface properties of an object are subtly changed during a saccade, below the threshold of detection, observers’ report their perception of the object as intermediate between its pre- and post-saccadic properties (Oostwoud Wijdenes, Marshall, & Bays, 2015). Furthermore, there is evidence this transsaccadic integration process follows principles of optimal probabilistic inference (e.g., Ernst & Banks, 2002), in that the perception reflects a weighted average, with weights that reflect the relative reliability of pre- and post-saccadic estimates of object features. Oostwoud Wijdenes et al. (2015) showed that when objects were viewed at different eccentricities before and after a saccade, the integrated percept was biased towards the feature value in the more foveal view. Similar shifts in perceived feature value were elicited when reliability was directly manipulated by adding external noise to the pre- or post-saccadic image. A number of studies (Ganmor, Landy, & Simoncelli, 2015; Hubner & Schutz, 2017; Stewart & Schütz, 2019a, 2019b; Wolf & Schutz, 2015) further demonstrated that a stable feature of an object visible both before and after a saccade can be identified more precisely than is possible from either the pre- or the post-saccadic view alone. This advantage is predicted as a result of averaging over independent noise in pre- and post-saccadic input. While the experimental demonstrations of this have typically involved items initially viewed in the periphery before a saccade brings them to the fovea, the behavioral benefits of averaging may be even more significant for items that remain peripheral, and therefore relatively poorly represented in the input, over the course of several fixations. For example, such peripheral-to-peripheral integration might allow an object to be identified that otherwise would not have been, whereas the benefits of peripheral-to-foveal integration will mostly be limited to resolving fine details.

However, our environment is rarely entirely static, and when changes occur during an eye movement, information from previous saccades may become actively misleading about the present state of the world. An apparent discrepancy between pre- and post-saccadic vision of an object could be the result of inherent variability in the visual system, in which case integration would be beneficial for accurate perception, or due to a real change in the object, in which case it would be harmful. The process of resolving this uncertainty has been termed causal inference (for a review, see Shams & Beierholm, 2010). In the context of transsaccadic integration of spatial information, a model formulated by Atsma, Maij, Koppen, Irwin, and Medendorp (2016) considers two possible causes of a discrepancy between pre- and post-saccadic estimates of object location: an inaccurate saccade or a change in the state of the world. In the former case, the goal is to integrate the two estimates, while in the latter, it is to segregate them and base decisions only on the post-saccadic input (Cicchini, Binda, Burr, & Morrone, 2013; Demeyer, De Graef, Wagemans, & Verfaillie, 2009; Stewart, Valsecchi, & Schütz, 2020). These possible causes and their resulting location estimates are then weighted, incorporating a prior belief favoring a stable world, so that larger intrasaccadic differences lead to responses weighted more towards the segregated post-saccadic estimate, while smaller differences lead to a percept weighted towards an integrated estimate.

Atsma et al.’s causal model makes inferences about, and on the basis of, a single visual parameter: object location. Real-world objects are made up of multiple features, and while internal variability may affect each feature independently, external causes will often change multiple features simultaneously. How should such cases be resolved? Given any evidence that an object has changed during an eye movement, the visual system might rely solely on the post-saccadic input; we describe such a process as *object-based*. Alternatively, evidence of a change in one feature could influence only the decision to integrate or segregate the pre- and post-saccadic estimates of that same feature, with independent decisions made for each object feature; we describe such a process as *feature-based*.

In this study, we investigated transsaccadic integration under circumstances involving changes to multiple features. We made intrasaccadic changes of varying magnitude to two features of a single object. We asked observers on every trial whether they had detected *any* change to the object, and also for an analogue estimate of one of the two features that they last perceived. Importantly, although we ask for a change detection response for any feature, since the differences in magnitude between the two features are uncorrelated, we can probabilistically infer which feature a change was detected in. This method revealed a correlation between detection of changes and transsaccadic integration within a feature dimension, but no evidence for interaction between feature dimensions.

## Methods

### Participants

36 participants took part in this study in total. All had normal or corrected-to-normal vision and were naïve as to the purpose of the experiment. The experiments were approved by the Cambridge Psychology Research Ethics Committee and informed consent was obtained in accordance with the Declaration of Helsinki.

### Apparatus and Stimuli

Stimuli were presented on a 27” Asus ROG PG279Q monitor (144 Hz refresh rate, 2560 × 1440 pixels), at a distance of 60 cm from the participant. The background of the screen was black (0.3 cd/m^2^) throughout the experiment. Eye position was tracked online at 1000 Hz with a desk-mounted EyeLink 1000 (SR Research). The stimuli were generated in Matlab using the Psychophysics Toolbox extensions (Kleiner, Brainard, & Pelli, 2007)

### Experiment 1

12 participants (5 male, 7 female, mean age 23.1, range 19–30) took part in this experiment. A summary of the procedure is shown in Figure 1a. Each trial began with the presentation of two dots (diameter, 0.5° of visual angle), located 6° horizontally left and right of the center of the screen. One dot was white (100 cd/m^2^) indicating the current location for fixation, while the other was gray (10 cd/m^2^) indicating the upcoming saccade target.

**Figure 1.**
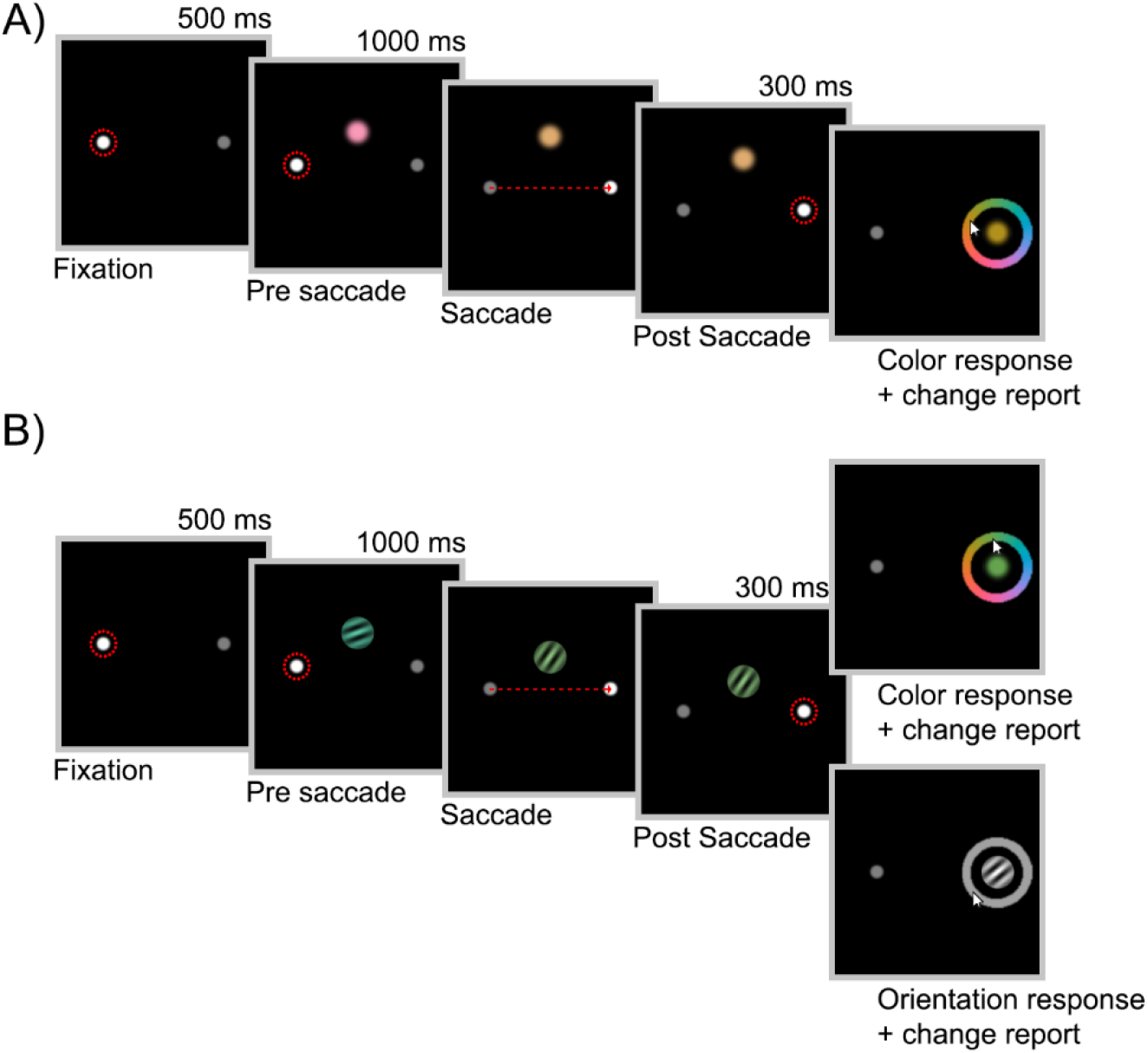
Experimental design. A) Trial sequence for Experiment 1. A color patch was presented that changed in color and/or location while the participant was making a saccade. They reported the last color they had seen and then whether they had detected any change in the stimulus. B) Experiment 2 followed the top path of this trial sequence (color and/or orientation change, followed by color report); Experiment 3 followed either the top or bottom path (color or orientation report) with equal frequency. Dashed red circles represent gaze fixations and dashed red arrows represent saccades. The stimulus changed as soon as gaze crossed the vertical midline of the screen.

Once the participant had maintained gaze within 1.5° of the fixation dot for 500 ms, a pre-saccadic stimulus appeared, consisting of a colored disk (diameter, 1°) located at one of four vertical displacements [–5°, –4°, +4°, +5°] from the screen center, randomly chosen with equal probability on each trial. The color of the pre-saccadic stimulus was randomly drawn from a circle in CIELAB space, centered on *L*\* = 74, *a*\* = *b*\* = 0, with a radius of 40. After 1000 ms, the two dots switched luminance, indicating that the participant should immediately make a saccade to the new fixation location on the opposite side of the screen. As soon as gaze position was detected to have crossed the vertical midpoint of the screen, the pre-saccadic stimulus was replaced with a post-saccadic stimulus.

The post-saccadic stimulus was identical to the pre-saccadic stimulus, except that its color was shifted clockwise or counterclockwise by 20°, 35° or 75° (equal frequency) on the color wheel, and its location was shifted vertically either up or down (equal probability) by 0°, 1.5° or 2.5° of visual angle (equal frequency). These values were selected based on pilot testing, to avoid floor or ceiling effects in the change detection responses.

After 300 ms of fixation at the new location, the post-saccadic stimulus was removed and a color wheel was displayed, centered on the fixation dot. The color wheel was randomly rotated from trial to trial. Participants were instructed to report the last color they had seen on the trial with a mouse click on the wheel. Moving the mouse cursor over the color wheel caused a disk to appear at fixation, with a color corresponding to the cursor location. After the color response, participants reported whether they had detected any change to the stimulus (regardless of whether the change was detected in color, location or both features) during the trial by clicking the left (no change) or right (change) mouse button. After the response, the next trial began with initial fixation at the opposite location to the previous trial. Note that the color estimation was always requested before the change detection response. This was intended to minimize any effect of the change detection response on the color estimate (e.g., a consistency bias that might have led participants to exaggerate the color difference from the pre-saccadic value after reporting a change).

If the participant made any erroneous saccades, took longer than 150 ms to initiate their saccade, finished their saccade further than 2.5° from the saccade target, or blinked before the response screen, the trial was immediately aborted. Aborted trials were presented again at the end of each block. Each block consisted of 54 successful trials. Participants performed either 6 blocks or 1.5 hours of the task, whichever took longer, resulting in a minimum of 324 successful trials and a maximum of 540.

### Experiment 2

12 participants (4 male, 7 female, 1 non-binary, mean age 23.8, range 18–31) took part in this experiment. The design was identical to Exp 1, with the following exceptions. The pre- and post-saccadic stimuli were colored Gabor patches, 1.5° in diameter, with a spatial frequency of 4.5 cycles/degree. The orientation of the pre-saccadic Gabor patch was drawn from a uniform distribution over all possible orientations. The orientation of the post-saccadic stimulus was rotated clockwise or counterclockwise by 10°, 30° or 90° (equal frequency) from that of the pre-saccadic stimulus. The colors of the pre- and post-saccadic stimuli varied as in Exp 1. The pre- and post-saccadic locations were the same on each trial. As in Exp 1, participants reported the last color they had seen and whether they had detected any change to the stimulus during the trial. Participants performed either 6 blocks or 1.5 hours of the task, whichever took longer, resulting in a minimum of 324 successful trials and a maximum of 540.

### Experiment 3

12 participants (5 male, 7 female, mean age 21.3, range 18–29) took part in this experiment. The design was identical to Exp 2, except that participants were unpredictably asked to report either color or orientation on each trial (equal frequency, randomly interleaved). On orientation report trials, a gray wheel appeared instead of the color wheel in the response stage of the task. This signaled the participant to report the last orientation they had seen with a mouse click at a corresponding point on the wheel. Moving the mouse cursor over the gray wheel prompted a grayscale Gabor patch to appear at fixation, with an orientation corresponding to the cursor position on the wheel. As in the previous experiments, participants also reported whether they had detected any change to the stimulus on each trial. Participants completed either 12 blocks or 3 hours of the task, whichever took longer, resulting in a minimum of 738 successful trials and a maximum of 900.

## Analysis

For each combination of participant, magnitude of color change and magnitude of location change (in Exp 1; orientation change in Exps 2 & 3), we calculated the mean proportion of trials on which a change was reported and a mean color bias. The latter was obtained by first rotating and reflecting the color response on each trial such that 0° corresponded to the pre-saccadic color, and positive values were in the direction of the post-saccadic color shift. The mean color bias was then calculated as the circular mean of these color values divided by the magnitude of the color change, such that 0 corresponded to the pre-saccadic color and 1 to the post-saccadic color. Orientation report trials in Experiment 3 were analyzed independently but identically to color report trials for the descriptive statistics, and in conjunction with the color report trials for the model analysis. Statistical tests of hypotheses were conducted using Bayesian ANOVAs and Bayesian *t*-tests in JASP (JASP Team, 2020) with default priors. The outcomes are reported as Bayes factors (BF), e.g., for a *t*-test, BF_10_ = 5 indicates that the strength of evidence for a difference is five times greater than for no difference; BF_01_ = 5 indicates the same strength of evidence favoring no difference. For the Bayesian ANOVA, models containing each of the main effects individually, a model containing both main effects and a null model incorporating only the intercept were compared using their Bayes Factors. For the sake of brevity, where the best fitting model contains both main effects, we report only the BF for this model versus the null. Otherwise, we report all relevant pairwise comparisons between models.

## Model

Although participants on each trial were asked only if they detected a change in either feature and to report their last perception of just one feature, which feature(s) they detected changes in and whether the responses were of integrated or segregated estimates can nonetheless be inferred using a probabilistic model. This is possible because the magnitudes of the changes in the two features were independent and controlled by the experimenter.

A pictorial representation of the model can be seen in Figure 2. For each trial in Experiment 1, the variables entered into the model were the pre-saccadic (−) and post-saccadic (+) feature values in color (*C*) and location (*L*) dimensions, 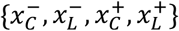, an analogue color estimate, 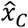, and a binary judgment of whether a change had occurred in either dimension, у ∈ {0,1}.

**Figure 2.**
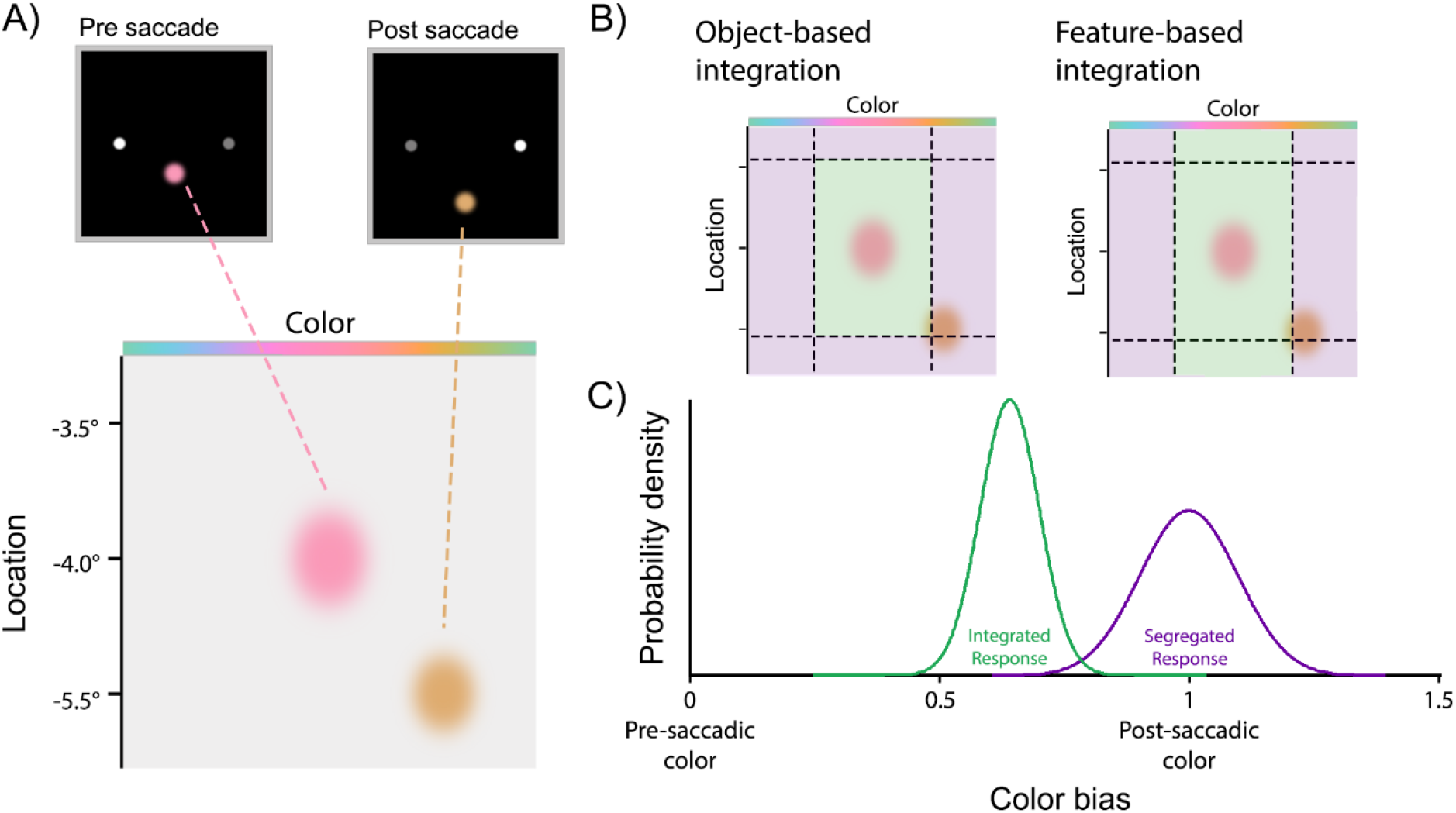
Pictorial representation of the optimal observer model. A) The pre- and post-saccadic color and location representations are corrupted by Gaussian noise, as represented by the bivariate Gaussians on the location and color feature spaces. B) Two-tailed thresholds, as represented by the dotted lines, are set for the purpose of detecting a change. If the post-saccadic stimulus exceeds a feature threshold, then a change is detected in the corresponding feature. For both models, if the post-saccadic representation lies in the green region, then pre- and post-saccadic representations will be integrated. If it falls in the purple region, then the representations will be kept separate. C) If the representations are integrated, then the analogue response will be centered on a point between the pre- and post-saccadic representations, as indicated by the green probability distribution. If the representations are segregated, then the analogue response will be centered on the post-saccadic representation (as indicated by the purple distribution) as the instruction was to report the last color perceived.

### Model specification

We assumed that internal representations of pre- and post-saccadic features were corrupted by independent Gaussian noise (wrapped onto the circular feature space in the case of color; Figure 2A),

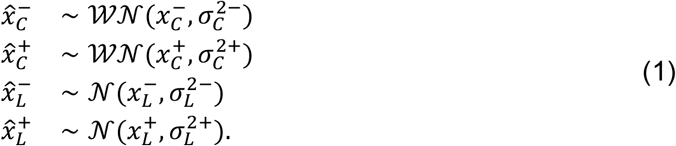

where 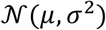 and 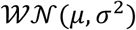 are the normal and wrapped normal distributions with mean μ and variance σ^2^. The observer detected a change if the estimated change magnitude in either dimension exceeded the threshold for that dimension (Figure 2B),

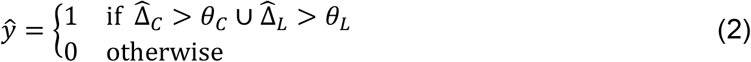

where 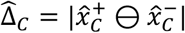, and 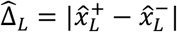 with ⊖ indicating subtraction on the circle. The observed detection response, у, reflected the internal detection state, ŷ, with a lapse rate, γ, i.e.,

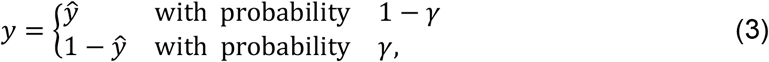

We compared predictions of two models that differed only in the conditions determining whether integration of pre- and post-saccadic features occurred in the reported dimension. In the *object-based integration* model, the conditions on integration matched those on detection, i.e., if the estimated change in either dimension exceeded threshold, the observer reported the internal post-saccadic estimate, otherwise they reported an integrated value based on both pre- and post-saccadic estimates (Figure 2C),

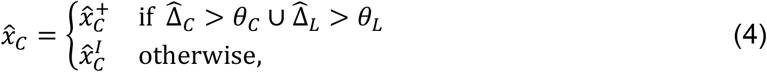

where the integrated estimate was a weighted average of pre- and post-saccadic estimates with weighting determined by their relative variances,

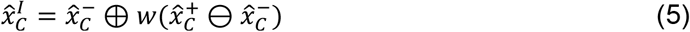

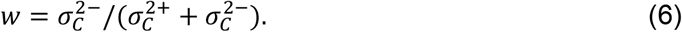

This weighting is optimal for combining normally distributed estimates (it is the minimum variance unbiased estimator) and so is also optimal for wrapped normal variables so long as their variances are not too large.

The *feature-based integration* model was identical, except that whether integration occurred in the reported dimension (color) depended only on whether a change was detected *in the same feature dimension*,

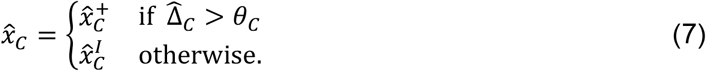

### Model fitting

The models are fully specified by the description above, however some further analysis is needed to determine likelihoods of model parameters. On a single trial, the probability of detecting a change in each dimension is

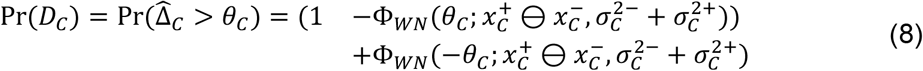

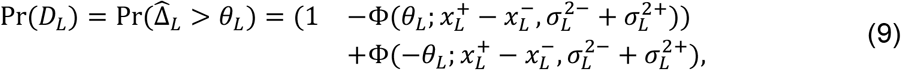

where 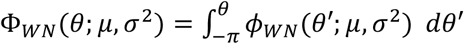 is the cumulative density function of the wrapped normal distribution, and Ф(θ μ, σ^2^) is the usual normal c.d.f. The joint probability of obtaining analogue response 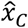 and not detecting a change is the same for both *object-based* and *feature-based* models:

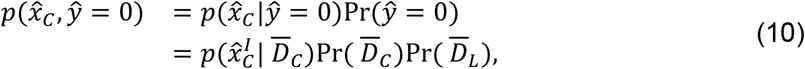

where 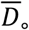 means no change detected in the indicated dimension, 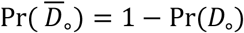, and noting that the distribution of the integrated color estimate is statistically independent of detection of a location change.

The joint probability of the analogue response and a change being detected differs between models. For the object-based model it is,

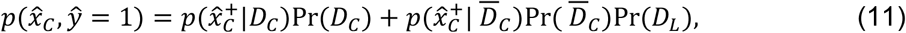

whereas for the feature-based model it is,

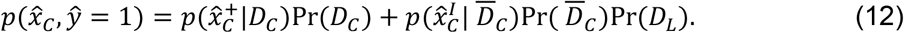

The conditional probability density for the analogue response given no color change was detected can be calculated as,

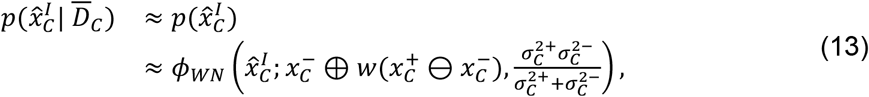

where, due to the circularity of the color space, both the independence of the integrated estimate from the detection state and its distribution as a wrapped normal with the stated variance are close approximations only. Using Bayes Theorem,

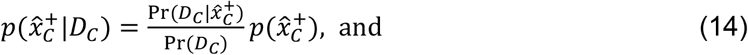

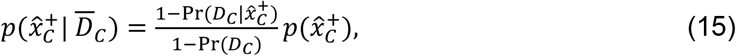

where

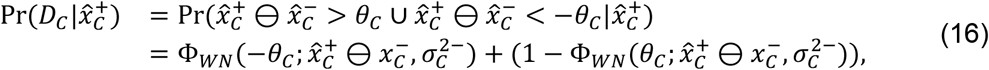
xs

Pr(*D_C_*) is given in Eq. 8, and 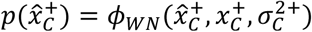. The probability of both responses is then given by

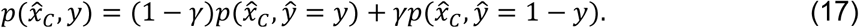

We used a non-linear optimization algorithm (*fminsearch* in MATLAB) to find parameters of the model that maximized this likelihood over all trials, for each participant separately. The pre- and post-saccadic variances in location, 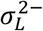 and 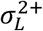 were not separable in the model so were fit with a single average parameter 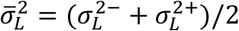. The models therefore each had six free parameters:

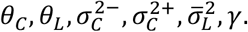

The models for Experiment 2 and the color-report trials of Experiment 3 were identical with the exception that orientation took the place of location and was treated as a circular variable, with individual estimates distributed as wrapped normals and addition and subtraction on the circle. The models for orientation-report trials of Experiment 3 incorporated these same changes, but with color interchanged with orientation. In this case pre- and post-saccadic orientation variances were separable and were treated as two free parameters, increasing the number of parameters in each model to seven.

All model comparisons were between models with the same number of free parameters, so we report the results as differences in log likelihood (ΔLL). Using BIC or AIC differences instead would have not changed conclusions, as these differences simply have magnitudes of twice the log likelihood difference.

## Results

In Experiment 1, we investigated whether changes in an object’s location would affect the transsaccadic integration of color information. Observers were instructed to report the color of the stimulus (the last color they saw if they believed it had changed) and whether they had detected any change to the color or location of the stimulus.

The effects of color and location change magnitude on the proportion of trials on which a change was reported are shown by the red symbols in Figures 3A and 3B respectively. As expected, detection probability increased with the size of the change in each feature. A two-way repeated-measures Bayesian ANOVA found that the best fitting model incorporated both main effects of color and location, BF_10_ = 6.48 ×10^17^. Follow-up Bayesian *t*-tests indicated that increasing changes in color led to increasing detection, with more change reports for 70° color differences than 35° (BF_10_ = 839.50) and for 35° than 20° (BF_10_ = 24.30). Similarly, there were more change reports for 2.5° location differences than 1.5° (BF_10_ = 25.57) and 1.5° than 0° (BF_10_ = 42.46).

**Figure 3.**
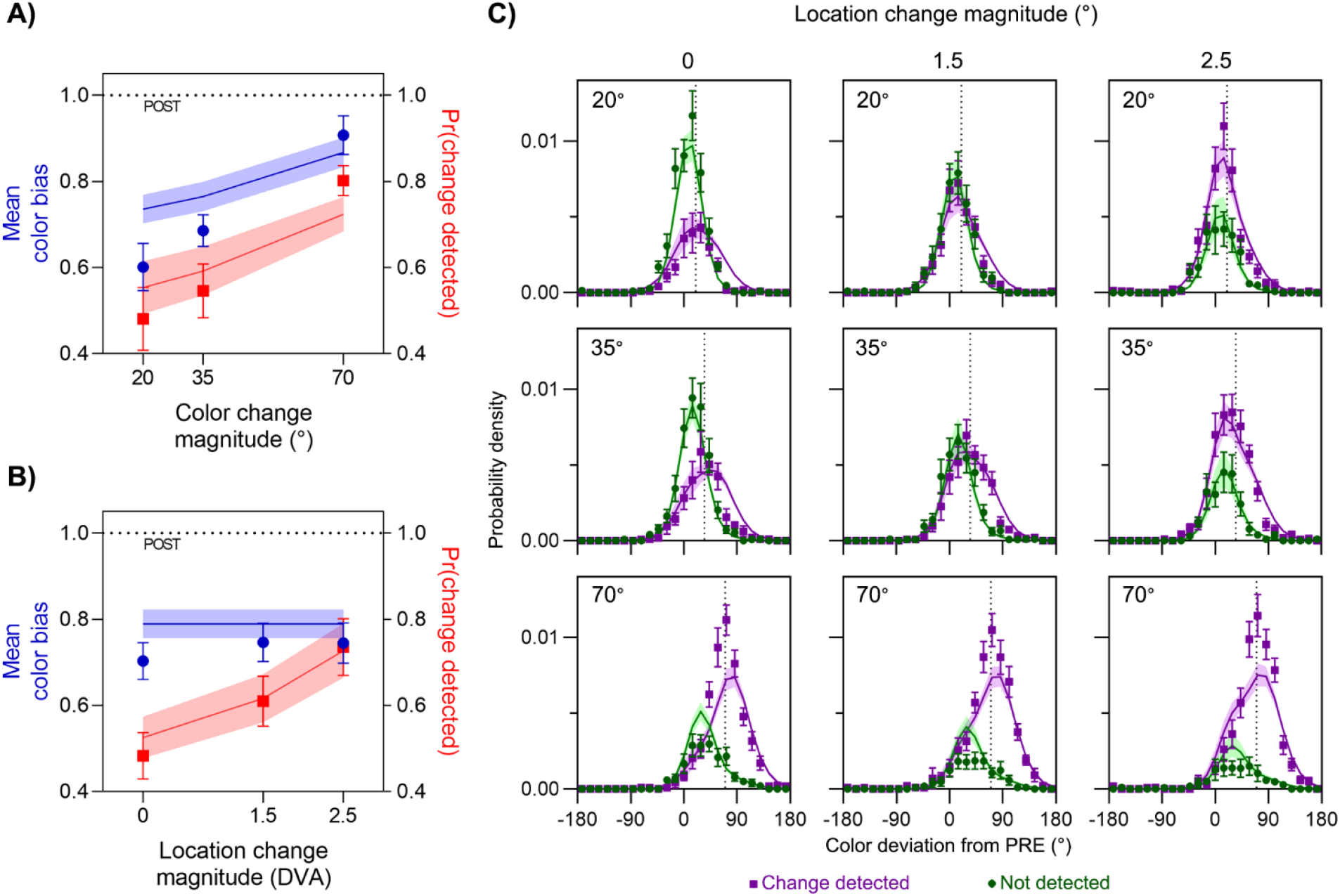
Results of Experiment 1. (A & B) Mean color bias (blue symbols, left y-axis) and mean frequency of detecting a change (red symbols, right y-axis) are plotted as a function of the magnitude of (A) color change and (B) location change. Curves show predictions of the best-fitting model, which was feature-based. (C) Distribution of reported color relative to pre-saccadic color (0°) and post-saccadic color (dotted vertical line in each panel), plotted separately for trials where a change was detected (purple) and not detected (green). Areas under each distribution reflect the frequency of detection. Each panel corresponds to a different pairing of color change (rows: magnitude indicated in top-left of each panel) and location change (columns: magnitude indicated at top). Error bars and error patches represent ± 1 S.E.

Having established that both the location and color manipulations were effective at modulating the probability of detecting a change to the stimulus, we examined their effects on its perceived color. We computed the bias in reported color as a value between 1 (the post-saccadic color) and 0 (the pre-saccadic color). Blue symbols in Figures 3A and 3B show the mean color bias for each color and location change magnitude. For the smallest color changes, despite the instruction to report only the last color seen, observers’ responses fell on average approximately mid-way (0.60 ± 0.05 [M ± SE]) between pre- and post-saccadic values. As the magnitude of color change increased, responses were increasingly biased towards the post-saccadic color, with a mean bias of 0.90 ± 0.05 for the largest color change.

In contrast, changes in location had minimal influence on transsaccadic integration of color information, as assessed by the mean color bias (Figure 3B, blue symbols), despite the clear effects on detection (red symbols). This qualitative difference was confirmed by a two-way Bayesian repeated-measures ANOVA, which supported the model incorporating only the main effect of color over the null hypothesis, BF_10_ = 3.39 × 10^13^, but showed the null hypothesis was favored over a model incorporating the main effect of location, BF_01_ = 7.04, and a model incorporating only the main effect of color was favored over a model which incorporated both main effects, BF = 3.93. Follow-up Bayesian *t*-tests indicated that color estimates were proportionally more biased towards the post-saccadic color as the color difference increased, as color estimates were more biased for 70° color differences than 35°, BF_10_ = 7.94 × 10^4^, and for 35° than 20°, BF_10_ = 2.30.

These results suggest that, while both color and location changes were detected by observers on some trials, and varying the change magnitude successfully modulated detectability over a large range for both features, only the detection of color changes had an influence on whether pre- and post-saccadic colors were integrated or segregated. To demonstrate the plausibility of this explanation and to create a direct test for it, we compared fits of two observer models to the joint distributions of color estimates and change reports (shown in Figure 3C).

In the *feature-based integration* model, observers reported an integrated color estimate (green distribution in Figure 2C) if the change in color across the saccade was not detected (green region in Figure 2B), and an estimate based only on the post-saccadic input if it was detected (purple region and distribution in Figures 2B and 2C respectively). This was contrasted with an *object-based integration* model, in which an integrated estimate was reported only if no change *in either feature* was detected. Model comparison favored the *feature-based* model (difference in log likelihood, ΔLL = 6.58 ± 3.23; 9 out of 12 participants). Predictions of this best-fitting model are shown as colored lines (with error patches) in each panel of Figure 3 (predictions for the object-based integration model can be seen in the Supplementary Materials).

This is strong evidence that the perception of a location change is disregarded when determining the perception of an object’s color after a saccade. We considered two explanations of this finding to have roughly equal plausibility. One possibility is that the decision to integrate or segregate features across a saccade is made independently in each different feature dimension (as assumed by our *feature-based integration* model). Alternatively, location changes specifically might be disregarded for purposes of integration. This could be because objects tend to change their locations more often than they change other features such color, making a change in location a less reliable marker of object continuity than other features. Indeed, changes of retinal location are an inevitable result of every saccade, and spatiotopic locations also change frequently because of physical interactions and gravity. To distinguish between these possibilities, we conducted Experiment 2.

### Experiment 2

This experiment was identical to Experiment 1, except that location changes were replaced with changes of stimulus orientation (illustrated by the top path in Figure 1B). The effects of color and orientation change magnitude on the proportion of trials on which a change was reported are shown by the red symbols in Figures 4A and 4B respectively. As expected, detection probability increased with the size of the change in each feature. A Bayesian ANOVA found that the best fitting model was one that incorporated both main effects of color and orientation, BF_10_ = 7.42 × 10^10^. Follow-up Bayesian *t*-tests indicated that increasing differences in color led to increasing detection, with more change reports for 70° color differences than 35° (BF_10_ = 31.24) and for 35° color differences than 20° (BF_10_ = 7.39). Weak evidence was found that greater differences in orientation also led to increasing detection, with more change reports for 35° orientation differences than 10° (BF_10_ = 2.34) and ambivalent evidence for any difference in change reports between 90° and 35° orientation differences (BF_01_ = 1.16).

**Figure 4.**
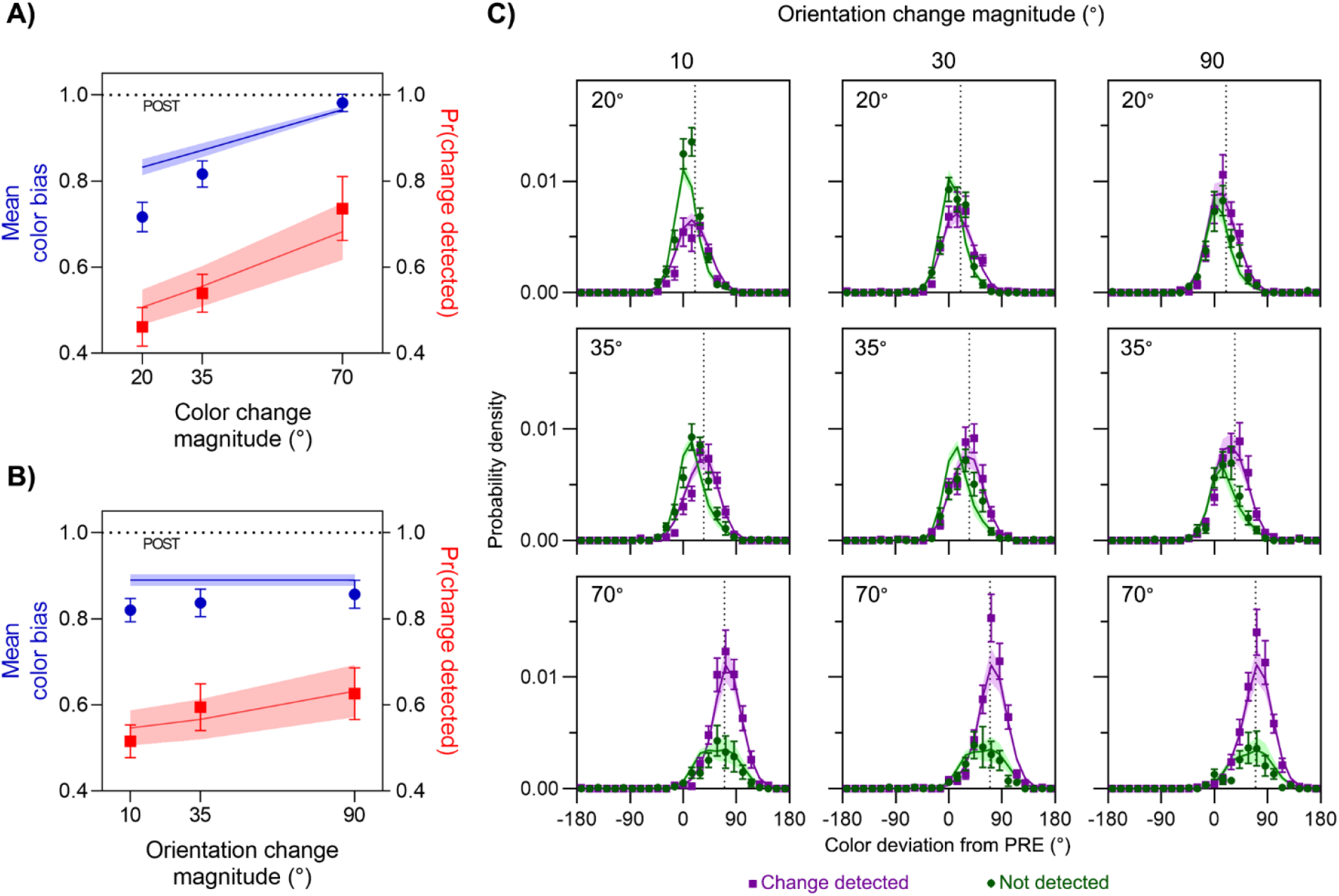
Results of Experiment 2. (A & B) Mean color bias (blue symbols, left y-axis) and mean frequency of detecting a change (red symbols, right y-axis) are plotted as a function of the magnitude of (A) color change and (B) orientation change. C) Distribution of reported color relative to pre-saccadic (0°) and post-saccadic color (dotted line in each panel), plotted separately for trials where a change was detected (purple) and not detected (green). Areas under each distribution reflect the frequency of detection. Each panel corresponds to a different pairing of color change (rows: magnitude indicated in top-left of each panel) and orientation change (columns: magnitude indicated at top).

Since both features were effective at modulating detection of changes in the stimulus, we examined their effects on perceived color. As shown by the blue symbols in Figure 4A, color responses were biased increasingly towards the post-saccadic value as the color change magnitude increased (from 0.72 ± 0.03 to 0.98 ± 0.02). However, changes in orientation had minimal effect on the integration of color information (blue symbols, Figure 4B), despite improving detection (red symbols). This was confirmed by a Bayesian ANOVA, which found that a model incorporating only the main effect of color was more likely than the null hypothesis, BF_10_ = 1.33 × 10^11^, but that the null hypothesis was favored over a model that incorporated only the main effect of orientation, BF_01_ = 8.18. The best fitting model incorporated only the main effect of color, being more likely than one that incorporated both main effects, BF = 6.26. Follow-up Bayesian *t*-tests indicated that color estimates were more biased towards the post-saccadic color as the color difference increased (70° vs 35°, BF_10_ = 1.47 × 10^4^; 35° vs 20°, BF_10_ = 118.92).

These results replicate those found for location changes in Experiment 1. While the detection of changes increased with the magnitude of change in both color and orientation changes, the integration of pre- and post-saccadic colors was affected only by the detection of color changes. We tested this by comparing fits of the *feature*- and *object*-*based integration* models to the joint distributions of color estimates and change reports (Figure 4C). Model comparison favored the *feature*-*based* model (ΔLL = 1.01 ± 0.36; 10 out of 12 participants).

This is evidence that the lower reliability of location as a marker of object correspondence is not the reason it is disregarded for the purpose of integrating color information across saccades. Instead, the evidence favors the explanation that the decision of whether or not to integrate a feature across a saccade depends only on the feature that is being integrated. However, we note that both Experiments 1 and 2 required an analogue report only in the color dimension, with the relevance of the other feature dimension limited to the binary change response. It is possible that this could have encouraged participants to pay more attention to color than location or orientation, which could influence feature binding (Treisman & Gelade, 1980). We conducted Experiment 3 to test this hypothesis.

### Experiment 3

In this experiment, we balanced attentional requirements by making unpredictable which feature participants would be asked to report on each trial. This is important as the amount of attention to the stimulus has been shown to affect perception in transsaccadic integration (Stewart & Schütz, 2018; Van der Stigchel, Schut, Fabius, & Van der Stoep, 2020). Half the trials were identical to those of Experiment 2. On the other half of trials, randomly interleaved, participants were asked to reproduce the last orientation they had seen, instead of the last color. This removed the asymmetry in task demands that might have caused participants to pay more attention to the color of the stimulus.

#### Color report trials

The effects of color and orientation change magnitude on the proportion of trials on which a change was reported are shown by the red symbols in Figures 5A and 5B, respectively. As in the previous two experiments, detection rates increased with magnitude of the change in each feature. A Bayesian ANOVA confirmed this, with the best model incorporating both main effects, BF_10_ = 1.16 × 10^17^. Follow-up Bayesian *t*-tests indicated that increasing differences in color led to more reports of changes (70° vs 35°, BF_10_ = 5.53 × 10^4^; 35° vs 20°, BF_10_ = 282.63) as did increasing differences in orientation (90° vs 35°, BF_10_ = 33.14; 35° vs 10°, BF_10_ = 19.73).

**Figure 5.**
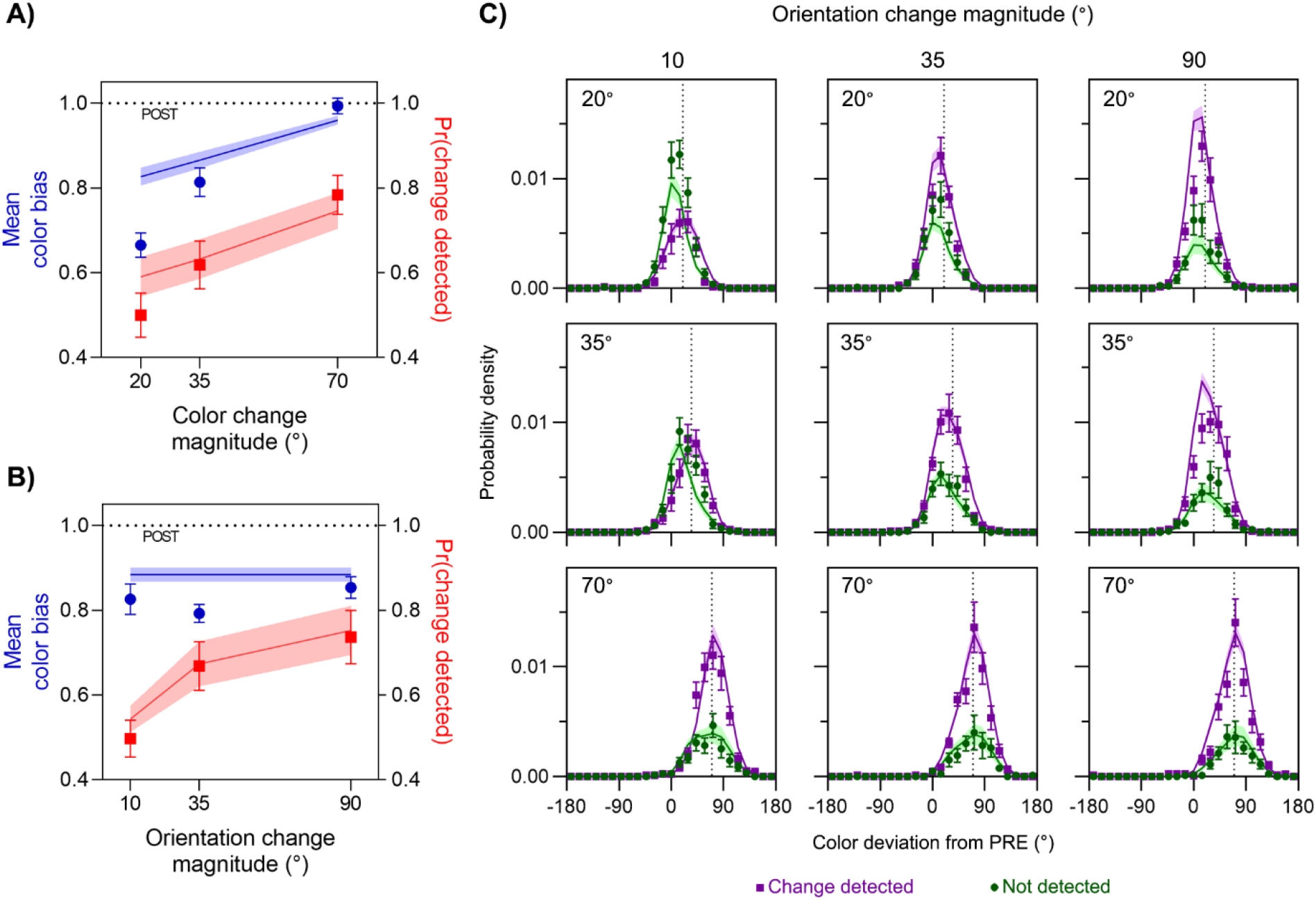
Results from the color-report trials in Experiment 3. (A & B) Mean color bias (blue symbols, left y-axis) and mean frequency of detecting a change (red symbols, right y-axis) are plotted as a function of the magnitude of (A) color change and (B) orientation change. C) Distribution of reported color relative to pre-saccadic (0°) and post-saccadic (dotted line in each panel) values, plotted separately for trials where a change was detected (purple) and not detected (green). Areas under each distribution reflect the frequency of detection. Each panel corresponds to a different pairing of color change (rows: magnitude indicated in top-left of each panel) and orientation change (columns: magnitude indicated at top).

Having established that both features were effective at modulating the probability of detecting a change in the stimulus, we turned to the effects on perceived color. As shown by the blue symbols in Figure 5A, color reports were increasingly biased towards the post-saccadic color (from 0.66 ± 0.03 to 0.99 ± 0.02) as the magnitude of the color change increased. However, as seen in blue symbols in Figure 5B, the magnitude of the orientation change had minimal effect on the color reports. This was confirmed by a Bayesian ANOVA, which found that the model incorporating only the main effect of color was favored over the null hypothesis, BF_10_ = 1.18 × 10^15^, whereas the null hypothesis was favored over the model incorporating only the main effect of orientation, BF_01_ = 5.24. Furthermore, the best fitting model incorporated only the main effect of color, being favored over a model incorporating both main effects, BF = 2.02. Follow-up Bayesian *t*-tests indicated that color estimates were proportionally more biased towards the post-saccadic color as the color difference increased (70° vs 35°, BF_10_ = 1.32 × 10^4^; 35° vs 20°, BF_10_ = 37.44).

These results indicate that, even when task demands were balanced across the two feature dimensions, the independence between color bias and orientation changes observed in Experiments 1 and 2 remained. This was confirmed by fitting the joint distributions of color estimates and change reports (shown in Figure 5C) with the *feature*-*based* and *object*-*based* integration models. Model comparison again favored the *feature*-*based* model (ΔLL = 3.65 ± 2.03; 10 out of 12 participants).

#### Orientation report trials

Although the choice of orientation changes in Experiment 3 was optimized for examining integration of color estimates, this experiment also gave us an opportunity to test if our conclusions generalized from color integration to integration of orientations.

As expected, all the effects of color and orientation change magnitude on detection that we had observed on color report trials were also present on orientation report trials (red symbols in Figures 6A and 6B). However, the effects of color and orientation changes on orientation estimates were the opposite of that seen for color estimates, in that orientation bias did not vary with the color change magnitude, but increased with increasing orientation change magnitude (from 0.77 ± 0.06 to 1.01 ± 0.01). This was confirmed by a Bayesian ANOVA which found that a model incorporating only the main effect of orientation was favored over the null hypothesis, BF_10_ = 164.46, and the null hypothesis was favored over the model incorporating only the main effect of color, BF_01_ = 8.10. The best fitting model incorporated only orientation, being favored over the model incorporating both main effects, BF = 7.63. Follow-up Bayesian *t*-tests indicated that orientation estimates were more biased towards the post-saccadic orientation as the orientation difference increased from 10° to 35° (BF_10_ = 20.58). There was no difference between 35° and 90° (BF_01_ = 2.53), presumably because the bias was already close to its ceiling value of one for 35° differences.

**Figure 6.**
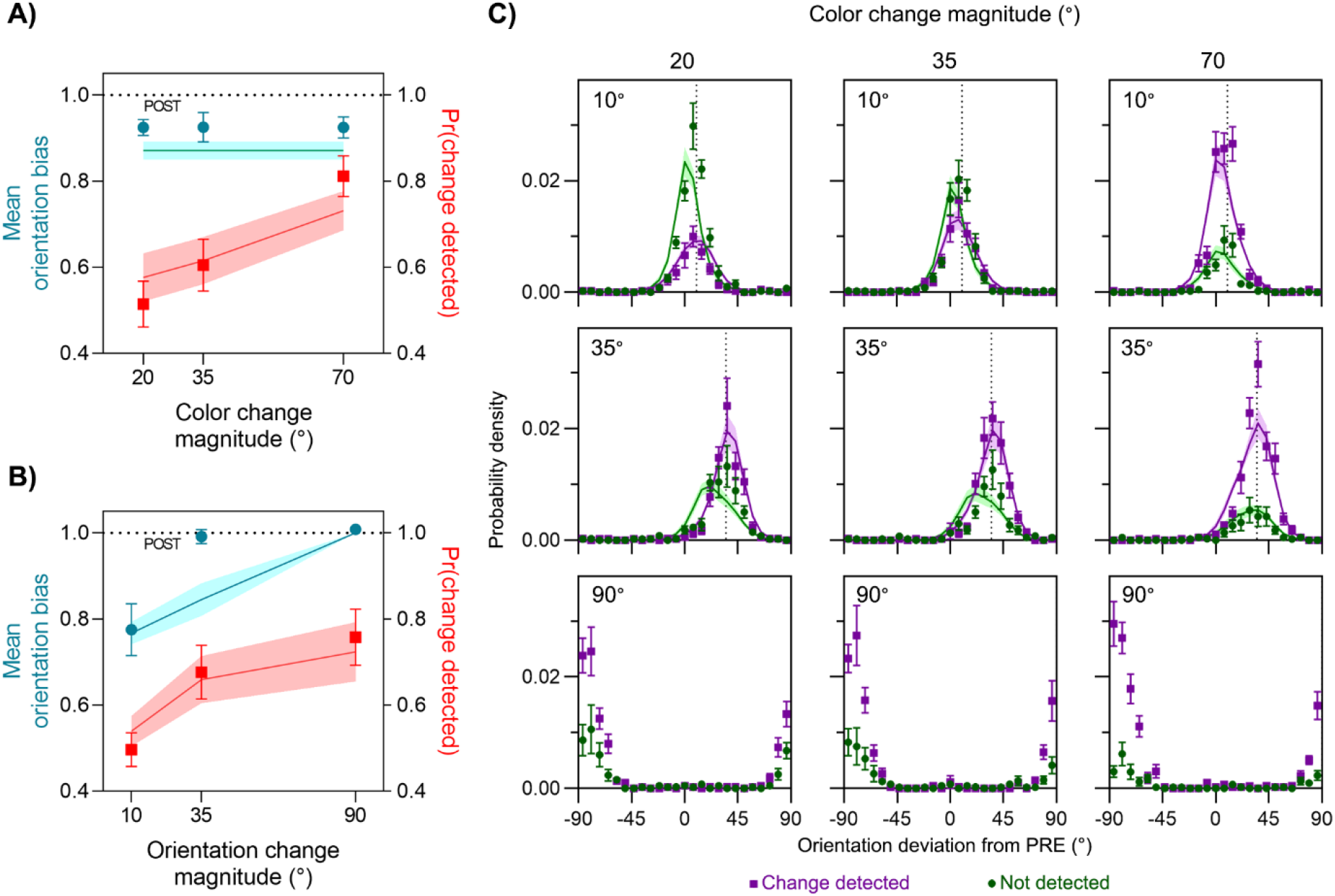
Results from the orientation-report trials in Experiment 3. (A & B) Mean orientation bias (cyan symbols, left y-axis) and mean frequency of detecting a change (red symbols, right y-axis) are plotted as a function of the magnitude of (A) color change and (B) orientation change. (C) Distribution of reported orientation relative to pre-saccadic (0°) and post-saccadic (dotted line in each panel) values, plotted separately for trials where a change was detected (purple) and not detected (green). Areas under each distribution reflect the frequency of detection. Each panel corresponds to a different pairing of orientation change (rows: magnitude indicated in top-left of each panel) and color change (columns: magnitude indicated at top). Model fits for the 90° orientation change condition are not available, as the optimal observer response for such a large change is not uniquely defined.

These results suggest that, while detection of color changes was used to determine whether or not to integrate color information across a saccade, changes in this feature were simultaneously ignored when determining whether or not to integrate orientation information (and vice versa). To test this directly, we fit the full data combining both color- and orientation-report trials with *feature*-*based* and *object*-*based* integration models. Parameters were shared across trial types such that, e.g., the same color variance parameters that contributed to change detection and analogue response probabilities on color-report trials also contributed to response probabilities on orientation-report trials. Note that trials with a 90° orientation change (i.e., when pre- and post-saccadic orientations were orthogonal) were omitted from the modelling, as the optimal integrated estimate is not clearly defined (Murray & Morgenstern, 2010). In practice, participants appear to have always reported the post-saccadic orientation on these trials (bottom panels of Figure 6C). Model comparison again favored the *feature*-*based* integration model (ΔLL = 1.29 ± 2.96; 8 out of 12 participants), in which integration in each feature dimension was determined independently of changes detected in the other feature dimension.

## Discussion

In everyday visual experience, objects that are otherwise stable tend not to abruptly change their properties during an eye movement. This makes integrating pre- and post-saccadic visual information a potentially advantageous strategy, as the combined estimate should be less influenced by perceptual noise. However, as the fidelity of pre-saccadic information available for transsaccadic computations is limited (Kong, Kroell, Aagten-Murphy, & Bays, in press), *any* evidence for a mismatch with the current environment could be considered justification to discount it and rely exclusively on the current (post-saccadic) input. However, across three experiments, we have found that the visual system operates according to different rules.

In Experiment 1, we investigated whether changes in an object’s location during a saccade altered the relative weighting given to pre- and post-saccadic colors in reports of the object’s color. We found no evidence for an influence of location changes on color integration, and our results were consistent with a model in which the decision to integrate or segregate color information was made solely on the basis of detecting color changes. In Experiments 2 and 3, we expanded this finding, by investigating whether the detection of changes in an object’s orientation would influence transsaccadic integration of its color, or vice versa. Experiment 3 also provided evidence against an alternative explanation in terms of unbalanced attention to the two features. In every case, changes in a particular feature only affected integration for that same feature. Note that while our experimental settings may have influenced participants to detect changes and segregate inputs more readily, as the frequency of intrasaccadic changes was much greater than usual for an object in the natural environment. However, while absolute rates of change detection may be inflated, this does not change our conclusions about the process of transsaccadic integration.

These results indicate that the integration of features held in pre-saccadic memory with post-saccadic visual input can occur in parallel for different features, separately from any overall decision about “same or different object”. In other words, an object may be seen to change in one feature at the same time that the perception of another feature reflects an averaging of pre- and post-saccadic values. For example, in trials with a large change in orientation and a small change in color, our results suggest that a participant who reported seeing a change would reproduce the post-saccadic orientation if asked for an orientation response but report an integrated pre- and post-saccadic color if asked for a color response. This pattern of responses may indicate that the visual system has adapted to take into account situations where objects change in one visual dimension while another remains stable, e.g., a ball rolling from light into shadow keeps a constant shape, but its luminance profile changes sharply. Whether a feature-based mechanism is advantageous in an ecological setting will depend on statistical regularities in the environment, in particular the frequency with which abrupt changes in different feature dimensions of an object co-occur. A truly optimal decision would also take into account the relative behavioral costs of incorrectly integrating features that have undergone a real change, versus incorrectly segregating features that could properly have been integrated.

An important point to address is the seeming discrepancy between a purely feature-based integration account, and the finding that changes in surface features aid in the detection of a displacement during a saccade (Tas et al., 2012). Our experiment was designed to test for feature interdependence in transsaccadic integration rather than change detection, meaning that we did not ask participants for separate reports of change detection in each feature dimension. Nonetheless, our results were quite accurately captured by a model in which detection occurred independently for each feature, and we found no evidence in any of the experiments for an effect of changes in the non-report feature on the analogue estimates of the report feature.

One possibility is that small dependencies between the two feature-level change detection processes were not detected in our study because they did not translate into object-level integration or segregation. However, there are also reasons why we might expect detection of intrasaccadic displacements, particularly ones parallel to the saccade direction as used by Tas et al. (2012), to be specifically sensitive to changes in other feature dimensions in a way that the features examined in the present study are not. First, unlike object colors or orientations, locations of objects in the visual field are necessarily altered by eye movements. Determining whether the object’s location in the world has changed requires (at least in the artificially sparse environments typical of Suppression of Saccadic Displacement studies) a comparison of its new retinal location to a prediction based on its pre-saccadic location and the eye movement vector. The latter is uncertain due to noise in the oculomotor system – especially so in the direction parallel to the saccade. This makes detection of transsaccadic location changes uniquely uncertain, and perhaps it is only in this case that feature changes in other dimensions have an appreciable influence.

Another possibility lies in the fact that our stimulus was distinct from and always some distance from the saccade target. While transsaccadic integration has been observed for objects at varying locations in the visual field (Oostwoud Wijdenes et al., 2015; Stewart & Schütz, 2019a), it is well established that the saccade target is automatically allocated attention (Deubel & Schneider, 2003) and stored in visual working memory (Bays & Husain, 2008; Schut, Van der Stoep, Postma, & Van der Stigchel, 2017). Based on the present results we cannot rule out the possibility that an object-based integration process operates only for the saccade target or in its vicinity. However, it does seem unlikely that this would be the case. A comparison between the results of Experiments 2 and 3 indicates that attention is not critical in determining whether the integration process is feature-based, and another recent study (Kong et al., in press) demonstrated that visual working memory is involved in the integration process for multiple stimuli, not just the saccade target.

A final reason why location changes might be treated differently by the visual system is to do with the statistics of objects in the natural environment, rather than saccades. If an object changing its location is in general less predictive of a change to its surface features than the converse, then an observer optimized for that environment will be more weakly influenced by location changes in judging whether a surface feature has changed, as compared to the influence of surface feature changes on judging location. This account would not predict differences between movements orthogonal and parallel to the saccade, and it would be interesting to know whether Tas et al.’s (2012) finding generalizes to orthogonal movements. The tendency for objects to move while their other properties remain stable may also underlie the privileged role of spatial location in binding other object features in memory (Golomb, Kupitz, & Thiemann, 2014; Kovacs & Harris, 2019; Pertzov & Husain, 2014; Schneegans & Bays, 2017).

Recently, the term *serial dependence* has been used to describe phenomena in which report of a visual stimulus is influenced by a statistically unrelated stimulus presented earlier, typically on the previous trial (J. Fischer & Whitney, 2014). Like transsaccadic integration (Irwin, 1991; Kong et al., in press; Oostwoud Wijdenes et al., 2015; Schut et al., 2017), serial dependence has been theorized to involve an integration of information obtained at two different points of time based on representations in working memory (Bliss, Sun, & D’Esposito, 2017; Fritsche, Mostert, & de Lange, 2017). Importantly for the purposes of this discussion, serial dependence also appears to happen on the level of features. One study found an attractive serial dependence in facial gender but a repulsive serial dependence of facial expression despite the two features being bound to the same face (Taubert, Alais, & Burr, 2016).

Note that there are also important differences between the two phenomena: transsaccadic integration involves saccadic suppression (e.g., Binda, Cicchini, Burr, & Morrone, 2009) and a translation of objects’ retinal locations; it is generally associated with a lack of awareness that a change has occurred; and effects sizes are generally greater than in serial dependence studies. For example, J. Fischer and Whitney (2014) found a peak bias of 8° in orientation towards the stimulus on the previous trial, which rapidly fell to zero as the difference between the previous and current stimuli increased. This result is actually something of an outlier in serial dependency studies, which have typically found even smaller effects, e.g., peak of 2–3° for Gabor patch orientation (Cicchini, Mikellidou, & Burr, 2017), 0.5– 2° for eye gaze direction (Alais, Kong, Palmer, & Clifford, 2018) and 2–3° for motion direction (C. Fischer et al., 2020). Notably, the findings of Wittenberg, Bremmer, and Wachtler (2008) – where small biases were observed towards a statistically unrelated stimulus separated from the current stimulus by a blank interval in addition to a saccade – might be more closely related to observations of serial dependence studies than transsaccadic integration.

In comparison, in Experiment 1, we found a peak effect of 11.0° in color bias in the 35° color change condition, which fell only as far as 7.9° at 20° color change, and 6.5° at 70° color change. If we were to confine ourselves to only those trials on which participants reported not detecting a change (which are the trials on which we believe integration took place), the effect sizes would be larger still: 9.5°, 17.9° and 32.6° respectively for 20°, 35° and 70° changes, i.e., approximately half the size of the color change. Recent work on serial dependence has found evidence to suggest that spatial location plays a distinct role as compared to other surface features (C. Fischer et al., 2020). However, further research is needed to determine whether and to what extent transsaccadic integration and serial dependence are expressions of a common mechanism of object correspondence.

## Acknowledgements

This research was supported by the Wellcome Trust (grant number 106926 to PMB). The funders had no role in study design, data collection and analysis, decision to publish, or preparation of the manuscript.

## Data Availability

Data and code associated with this study will be made publicly available on publication.

## References

Aagten-Murphy, D., & Bays, P. M. (2018). Functions of Memory Across Saccadic Eye Movements. Curr Top Behav Neurosci. doi:10.1007/7854_2018_66

Alais, D., Kong, G., Palmer, C., & Clifford, C. (2018). Eye gaze direction shows a positive serial dependency. Journal of vision, 18(4), 11. doi:10.1167/18.4.11

Atsma, J., Maij, F., Koppen, M., Irwin, D. E., & Medendorp, W. P. (2016). Causal Inference for Spatial Constancy across Saccades. PLoS Comput Biol, 12(3), e1004766. doi:10.1371/journal.pcbi.1004766

Bays, P. M., & Husain, M. (2008). Dynamic shifts of limited working memory resources in human vision. Science, 321(5890), 851–854. doi:10.1126/science.1158023

Binda, P., Cicchini, G. M., Burr, D. C., & Morrone, M. C. (2009). Spatiotemporal Distortions of Visual Perception at the Time of Saccades. The Journal of Neuroscience, 29(42), 13147–13157. doi:10.1523/jneurosci.3723-09.2009

Bliss, D. P., Sun, J. J., & D’Esposito, M. (2017). Serial dependence is absent at the time of perception but increases in visual working memory. Scientific Reports, 7(1), 14739–14739. doi:10.1038/s41598-017-15199-7

Bridgeman, B., Hendry, D., & Stark, L. (1975). Failure to detect displacement of the visual world during saccadic eye movements. Vision Res, 15(6), 719–722. doi:10.1016/0042-6989(75)90290-4

Burr, D. C., & Morrone, M. C. (2011). Spatiotopic coding and remapping in humans. Philos Trans R Soc Lond B Biol Sci, 366(1564), 504–515. doi:10.1098/rstb.2010.0244

Cavanagh, P., Hunt, A. R., Afraz, A., & Rolfs, M. (2010). Visual stability based on remapping of attention pointers. Trends Cogn Sci, 14(4), 147–153. doi:10.1016/j.tics.2010.01.007

Cicchini, G. M., Binda, P., Burr, D. C., & Morrone, M. C. (2013). Transient spatiotopic integration across saccadic eye movements mediates visual stability. J Neurophysiol, 109(4), 1117–1125. doi:10.1152/jn.00478.2012

Cicchini, G. M., Mikellidou, K., & Burr, D. (2017). Serial dependencies act directly on perception. Journal of vision, 17(14), 6–6. doi:10.1167/17.14.6

Demeyer, M., De Graef, P., Wagemans, J., & Verfaillie, K. (2009). Transsaccadic identification of highly similar artificial shapes. Journal of vision, 9(4), 28–28. doi:10.1167/9.4.28

Demeyer, M., De Graef, P., Wagemans, J., & Verfaillie, K. (2010). Object form discontinuity facilitates displacement discrimination across saccades. Journal of vision, 10(6), 17–17. doi:10.1167/10.6.17

Deubel, H., & Schneider, W. X. (2003). Delayed Saccades, but Not Delayed Manual Aiming Movements, Require Visual Attention Shifts. Annals of the New York Academy of Sciences, 1004(1), 289–296. doi:https://doi.org/10.1196/annals.1303.026

Ernst, M. O., & Banks, M. S. (2002). Humans integrate visual and haptic information in a statistically optimal fashion. Nature, 415(6870), 429–433. doi:10.1038/415429a

Fischer, C., Czoschke, S., Peters, B., Rahm, B., Kaiser, J., & Bledowski, C. (2020). Context information supports serial dependence of multiple visual objects across memory episodes. Nature Communications, 11(1), 1932. doi:10.1038/s41467-020-15874-w

Fischer, J., & Whitney, D. (2014). Serial dependence in visual perception. Nature Neuroscience, 17, 738. doi:10.1038/nn.3689

Fritsche, M., Mostert, P., & de Lange, F. P. (2017). Opposite Effects of Recent History on Perception and Decision. Current Biology, 27(4), 590–595. doi:https://doi.org/10.1016/j.cub.2017.01.006

Ganmor, E., Landy, M. S., & Simoncelli, E. P. (2015). Near-optimal integration of orientation information across saccades. Journal of vision, 15(16), 8. doi:10.1167/15.16.8

Golomb, J. D., Kupitz, C. N., & Thiemann, C. T. (2014). The influence of object location on identity: a “spatial congruency bias”. J Exp Psychol Gen, 143(6), 2262–2278. doi:10.1037/xge0000017

Hubner, C., & Schutz, A. C. (2017). Numerosity estimation benefits from transsaccadic information integration. Journal of vision, 17(13), 12. doi:10.1167/17.13.12

Irwin, D. E. (1991). Information integration across saccadic eye movements. Cognitive psychology, 23(3), 420–456. doi:10.1016/0010-0285(91)90015-g

JASP Team. (2020). JASP (Version 0.12)[Computer software]. In. Retrieved from https://jasp-stats.org/.

Kleiner, M., Brainard, D., & Pelli, D. (2007). What’s new in Psychtoolbox-3. Perception, 36 ECVP Abstract Supplement.

Kong, G., Kroell, L. M., Aagten-Murphy, D., & Bays, P. M. (in press). Resource Limitations in transsacadic integration. Journal of vision.

Kovacs, O., & Harris, I. M. (2019). The role of location in visual feature binding. Atten Percept Psychophys. doi:10.3758/s13414-018-01638-8

Melcher, D., & Morrone, M. C. (2015). Nonretinotopic visual processing in the brain. Vis Neurosci, 32, E017. doi:10.1017/s095252381500019x

Murray, R. F., & Morgenstern, Y. (2010). Cue combination on the circle and the sphere. Journal of vision, 10(11), 15–15.

Niemeier, M., Crawford, J. D., & Tweed, D. B. (2003). Optimal transsaccadic integration explains distorted spatial perception. Nature, 422(6927), 76–80.

Oostwoud Wijdenes, L., Marshall, L., & Bays, P. M. (2015). Evidence for Optimal Integration of Visual Feature Representations across Saccades. The Journal of Neuroscience, 35(28), 10146–10153. doi:10.1523/jneurosci.1040-15.2015

Pertzov, Y., & Husain, M. (2014). The privileged role of location in visual working memory. Atten Percept Psychophys, 76(7), 1914–1924. doi:10.3758/s13414-013-0541-y

Poth, C. H., Herwig, A., & Schneider, W. X. (2015). Breaking Object Correspondence Across Saccadic Eye Movements Deteriorates Object Recognition. Front Syst Neurosci, 9(176). doi:10.3389/fnsys.2015.00176

Rensink, R. A., O’Regan, J. K., & Clark, J. J. (1997). To See or not to See: The Need for Attention to Perceive Changes in Scenes. Psychological Science, 8(5), 368–373. doi:10.1111/j.1467-9280.1997.tb00427.x

Schneegans, S., & Bays, P. M. (2017). Neural Architecture for Feature Binding in Visual Working Memory. The Journal of neuroscience : the official journal of the Society for Neuroscience, 37(14), 3913–3925. doi:10.1523/JNEUROSCI.3493-16.2017

Schut, M. J., Van der Stoep, N., Postma, A., & Van der Stigchel, S. (2017). The cost of making an eye movement: A direct link between visual working memory and saccade execution. Journal of vision, 17(6), 15–15. doi:10.1167/17.6.15

Shams, L., & Beierholm, U. R. (2010). Causal inference in perception. Trends Cogn Sci, 14(9), 425–432. doi:10.1016/j.tics.2010.07.001

Stewart, E. E. M., & Schütz, A. C. (2018). Attention modulates trans-saccadic integration. Vision Res, 142, 1–10. doi:https://doi.org/10.1016/j.visres.2017.11.006

Stewart, E. E. M., & Schütz, A. C. (2019a). Transsaccadic integration benefits are not limited to the saccade target. J Neurophysiol, 122(4), 1491–1501. doi:10.1152/jn.00420.2019

Stewart, E. E. M., & Schütz, A. C. (2019b). Transsaccadic integration is dominated by early, independent noise. Journal of vision, 19(6), 17–17. doi:10.1167/19.6.17

Stewart, E. E. M., Valsecchi, M., & Schütz, A. C. (2020). A review of interactions between peripheral and foveal vision. Journal of vision, 20(12), 2–2. doi:10.1167/jov.20.12.2

Tas, A. C., Moore, C. M., & Hollingworth, A. (2012). An object-mediated updating account of insensitivity to transsaccadic change. Journal of vision, 12(11), 18–18. doi:10.1167/12.11.18

Taubert, J., Alais, D., & Burr, D. (2016). Different coding strategies for the perception of stable and changeable facial attributes. Scientific Reports, 6(1), 32239. doi:10.1038/srep32239

Treisman, A., & Gelade, G. (1980). A feature-integration theory of attention. Cognitive psychology, 12(1), 97–136.

Van der Stigchel, S., Schut, M. J., Fabius, J., & Van der Stoep, N. (2020). Transsaccadic perception is affected by saccade landing point deviations after saccadic adaptation. Journal of vision, 20(9), 8–8. doi:10.1167/jov.20.9.8

Wexler, M., & Collins, T. (2014). Orthogonal steps relieve saccadic suppression. Journal of vision, 14(2), 13–13. doi:10.1167/14.2.13

Wittenberg, M., Bremmer, F., & Wachtler, T. (2008). Perceptual evidence for saccadic updating of color stimuli. Journal of vision, 8(14), 9.1–9. doi:10.1167/8.14.9

Wolf, C., & Schutz, A. C. (2015). Trans-saccadic integration of peripheral and foveal feature information is close to optimal. Journal of vision, 15(16), 1. doi:10.1167/15.16.1

